# Chromatin structure around long non-coding RNA (lncRNA) genes in *Schistosoma mansoni* gonads

**DOI:** 10.1101/2024.02.22.581540

**Authors:** Ronaldo de Carvalho Augusto, Thomas Quack, Christoph G. Grevelding, Christoph Grunau

## Abstract

In this study, we employed a total of eight distinct modifications of histone proteins (H3K23ac, H3K27me3, H3K36me3, H3K4me3, H3K9ac, H3K9me3, H4K12ac, and H4K20me1) to discern the various chromatin colors encompassing lncRNA genes in both mature and immature gonads of the human parasite *Schistosoma mansoni*. Our investigation revealed that these chromatin colors exhibit a tendency to aggregate based on the similarities in their metagene shapes, leading to the formation of less than six distinct clusters. Moreover, these clusters can be further grouped according to their resemblances by shape, which are co-linear with specific regions of the genes and potentially associated with transcriptional stages.

## Introduction

Schistosomiasis, a parasitic disease caused by members of the *Schistosoma* genus, represents a significant public health burden in tropical and subtropical regions, affecting 250 million people worldwide. The life cycle of these parasitic flatworms involves complex interactions with both intermediate and definitive hosts, resulting in chronic infections that can lead to severe morbidity and mortality if left untreated. Despite its substantial impact on global health, schistosomiasis has received relatively limited attention in comparison to other neglected tropical diseases, and the intricacies of its molecular mechanisms remain incompletely understood (McManus et al., 2018).

Among the platyhelminths schistosomes are exceptional in evolution because they are the only representatives, which have evolved separate sexes. Furthermore, schistosomes exhibit a nearly unique sexual biology because the maturation of the female gonad depends on a permanent pairing-contact with a male partner (Grevelding, 2004; Kunz, 2001; Loverde and Chen, 1991; Popiel and Basch, 1984). Among others, pairing induces the final differentiation of the ovary, which in an unpaired females consists of stem cell-like, undifferentiated oogonia that do not enter meiosis. After pairing, however, mitoses of gonadal cells are induced that finally lead to the differentiation of oocytes entering meiosis (Den Hollander and Erasmus, 1985; Fitzpatrick and Hoffmann, 2006). These and other changes are accompanied by a remarkable growth of the paired female, and they are mediated by complex signalling systems as well as many genes expressed in a pairing-dependent manner, including gonadal genes. Although in the male no morphological changes are observed following pairing, previous transcriptomics data of protein-coding genes of whole worms (and their gonads) revealed an influence of the pairing status also on gene expression in males, including their testes (Lu et al., 2016; Lu et al., 2019).

Recent advances in genomics and transcriptomics have shed light on the role of long non-coding RNAs (lncRNAs) in the regulation of gene expression and the modulation of biological processes across various organisms. LncRNAs, a class of non-protein-coding transcripts with a length exceeding 200 nucleotides, have emerged as crucial players in diverse cellular and physiological functions, including development, 1 differentiation, and disease pathogenesis (Ponting et al., 2009). Although the role of lncRNAs has been extensively investigated in various organisms, their involvement in schistosome biology remains largely unexplored territory.

Understanding the regulatory mechanisms governing schistosome lncRNA gene expression would allow for elucidating key aspects of their biology, including host-parasite interactions, immune evasion, and developmental transitions. In this context, the exploration of lncRNAs in schistosomes presents a promising avenue for uncovering novel regulatory elements and mechanisms that may contribute to the parasites’ adaptation and survival within their hosts.

In *Schistosoma mansoni*, lncRNA have been shown to affect pairing status, survival, and reproduction in adult worms, making them potential therapeutic targets (Silveira et al., 2023). Additionally, lncRNAs have been identified in different developmental stages of *S. mansoni*, suggesting their involvement in stage-specific gene expression and potential as biomarkers (Rocha et al., 2022). Furthermore, the expression patterns of lncRNAs in *S. mansoni* have been characterized at the single-cell level, revealing tissue-specific expression and regulated expression programs (Morales-Vicente et al., 2022). In *Schistosoma japonicum* infection, lncRNAs have been implicated in the formation of liver granulomas and the polarization of macrophages, providing a potential target for treatment (Zhao et al., 2023).

Intergenic, antisense, and sense lncRNAs are differentially expressed after treatment 5-AzaC, an inhibitor of DNA/RNA methylases, in *S. mansoni* females. These differentially expressed lncRNAs belong to co-expression network modules related to male metabolism, and they are also differentially expressed in unpaired females compared with paired females and their ovaries, respectively (Amaral et al., 2020). Re-analysis of single-cell RNA-sequencing data from adult *S. mansoni* identified the expression patterns of lncRNAs in different cell types including specific lncRNAs in the gonads (Rocha et al., 2022).

Given the stage- and cell specific expression of lncRNA and its sensitivity to an epigenetic stressor (Amaral et al., 2020), we hypothesized that lncRNA gene function could be influenced by chromatin structure around their genes. Chromatin, the complex of DNA and proteins that constitutes the structural framework of eukaryotic genomes, plays a pivotal role in controlling gene function and maintaining genome integrity. Over the past few decades, extensive research has revealed the remarkable diversity and complexity of chromatin organization, leading to the emergence of the concept of “chromatin colors” or “chromatin flavors.” These metaphoric terms describe, in analogy to the additive primary colors that can produce the full spectrum of visible colors, the idea that individual compositions of the chromatin can be combined into different functional states (“colors”) that it can adopt to orchestrate gene regulation and cellular identity (Strahl and Allis, 2000).

We used here 8 different posttranslational histone modifications (H3K23ac, H3K27me3, H3K36me3, H3K4me3, H3K9ac, H3K9me3, H4K12ac, and H4K20me1) to identify the chromatin colors surrounding lncRNA genes in gonads, ovaries and testes, isolated from paired and unpaired *S. mansoni*. We found that chromatin colors cluster by similarity of their metagene shapes into less than 6 clusters, which can be grouped into similar shapes that appear to be associated with parts of the genes, the pairing status of the adults from which the gonads originated, and potentially with transcription steps.

## Results

### 500bp is the optimal bin size for ChromstaR analysis

ChromstaR (Taudt et al., 2016) is a R-package for the analysis of ChIP-Seq data based on Hidden-Markov-Model (HMM). Regions of the genome that are enriched in specific histone modifications (“peaks”) are identified based on transition probabilities. ChromstaR allows to combine individual histone modifications into combinational states (“chromatin colours”). It requires to establish empirically a bin size prior to the full analysis. For this we used comparison with other peakcalling methods and visual inspection under IGV for the relatively sharp H3K4me3 peaks. Best concordance of peak detection between MACS2, Peakranger, and ChromstaR was obtained at a bin size of 500 bp with input BAM as reference. Visual inspection of peakcalling using MACS bedgraph, and Peakranger, and ChromstaR wig files, and BED files from ChromstaR revealed excellent concordance of peak representation and peak detection by ChromstaR with these parameters. Metagene profiles were similar for the tree methods and showed high similarity between replicates. The modified lines in the univariate fits of ChromstaR were flat, an empirical indicator for good model fitting. In conclusion, further analyses were done with binsize 500 bp and input BAM as references.

### There are sex-specific differences in chromatin profiles around lncRNA genes in gonads of S. mansoni

Based on a previously established organs isolation method (Hahnel et al., 2013), we used pairing-experienced females (bisex females, bF) and males (bM) as well as used pairing-unexperienced females (sF, single sex female) and males (sM) for isolation of their gonads, ovaries from bF (bO) and sF (sO) and testes of bM (bT) and sM (sT). Next, we performed ChIP-Seq with the abovementioned 8 antibodies. We aligned ChIP-seq reads to the genome and generated metagene profiles *i*.*e*. log(observed/expected) of color enrichment spanning 2 kb upstream of the transcription start site (TSS) to 2 kb downstream of the transcription end site (TES). We noticed that some profiles had similar shapes and used unsupervised clustering to sort similar profiles into clusters. We allowed up to 6 clusters but considered only clusters with at least 4 colors. Cluster 0 contains the “flatliners” *i*.*e*., no enrichment around lncRNA genes. Number of profiles in clusters are listed in table 1. By visual inspection of cumulated log(obs/exp) metagene profiles, we realized that these profiles can be grouped by their shapes (figure 1). These groups were different in both sexes and varied also between mature and immature gonads. We tentatively assigned the different cluster groups to parts of genes. It should be notes that clusters were independently numbered *e*.*g*. cluster 1 in bT has not the same composition as cluster 1 in sT. For the composition of the clusters, contact the authors.

**Table 1:** Numbers of testes from single-sex males (sT; extracted from worms representing clonal, unisexual worm populations), bisex testes (bT; extracted from worms representing clonal, bisexual worm populations), single-sex ovaries (sO; extracted form ss females) and bi-sex ovaries (bO; extracted from bs females). In bold given are clusters that were used for further analyses.

| cluster | sT | bT | sO | bO |
| --- | --- | --- | --- | --- |
| 0 (flatline) | 19 | 58 | 45 | 37 |
| 1 | **57** | **160** | **163** | **140** |
| 2 | **162** | **13** | **41** | **68** |
| 3 | **6** | **18** | 2 | **5** |
| 4 | 2 | 1 | 1 | 1 |
| 5 | **6** | **5** | 1 | 1 |
| 6 | 4 | 1 | 1 | 2 |

**Table 2:** Assignment of clusters to part of genes (see also figure 1) Abbreviations as in the main text.

| Part of gene | sT | bT | sO | bO |
| --- | --- | --- | --- | --- |
| TSS | Cluster 3 | Cluster 1 |  | Cluster 3 |
| Gene body | Cluster 6, 2 | Cluster 3, 2 | Cluster 1 | Cluster 2 |
| TES and downstream | Cluster 1, 5 | Cluster 5 |  |  |
| Absent in gene body |  |  | Cluster 2 | Cluster 1 |

**Fig. 1a.**
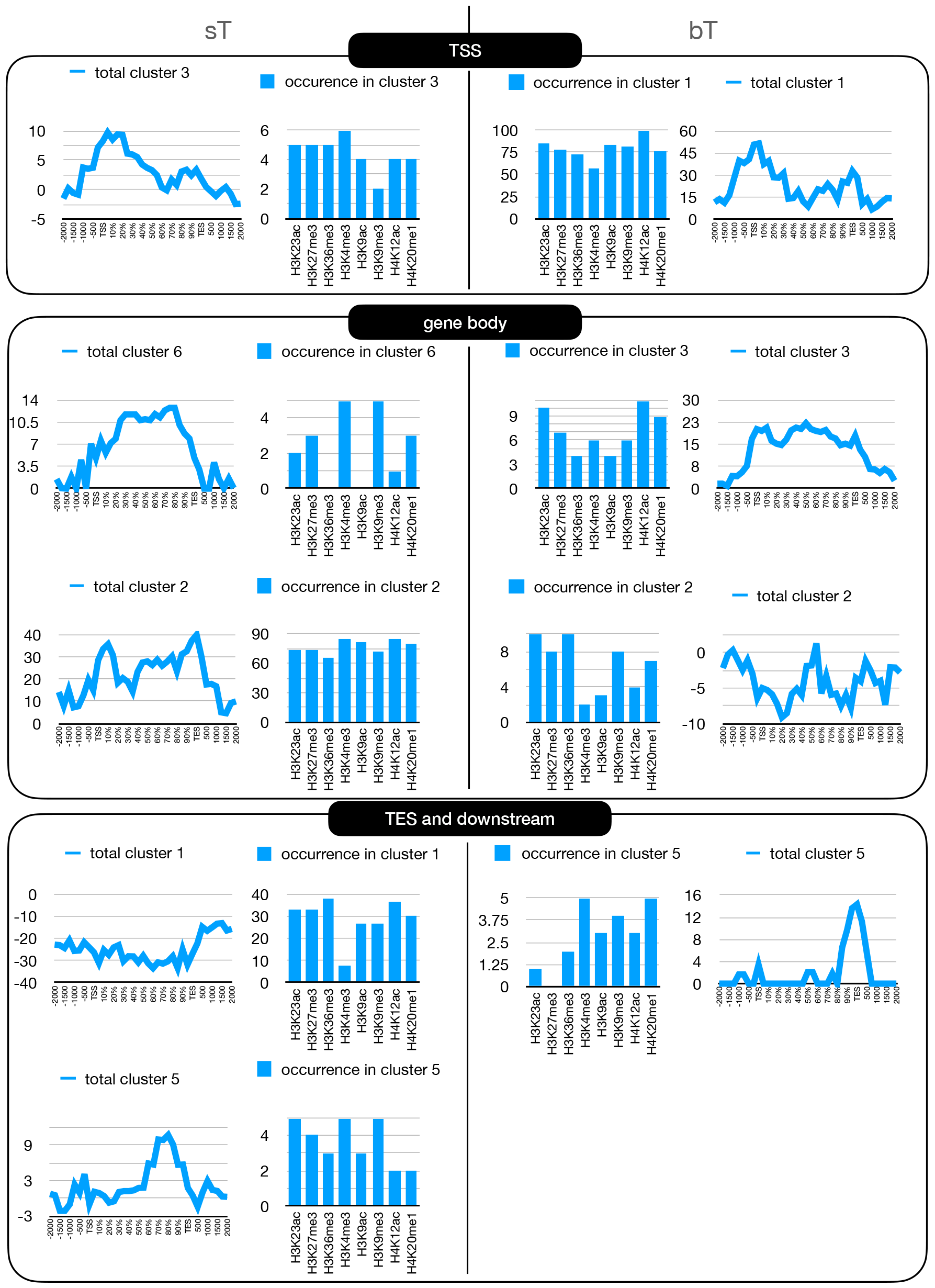
Metagene profiles of testes of single-sex, pairing-unexperienced males (sT), and bisex, pairing-experienced males (bT), combined into clusters by shape and associated with gene parts. X-axis 2 kb upstream of TSS and 2 kb downstream of TES. Y-axis log(obs/exp) totaled over all profiles in the cluster. Histograms show the absolute number of occurrences of a histone modification in the chromatin colors. Squares surround groups of profiles with similar shape.

**Fig. 1b.**
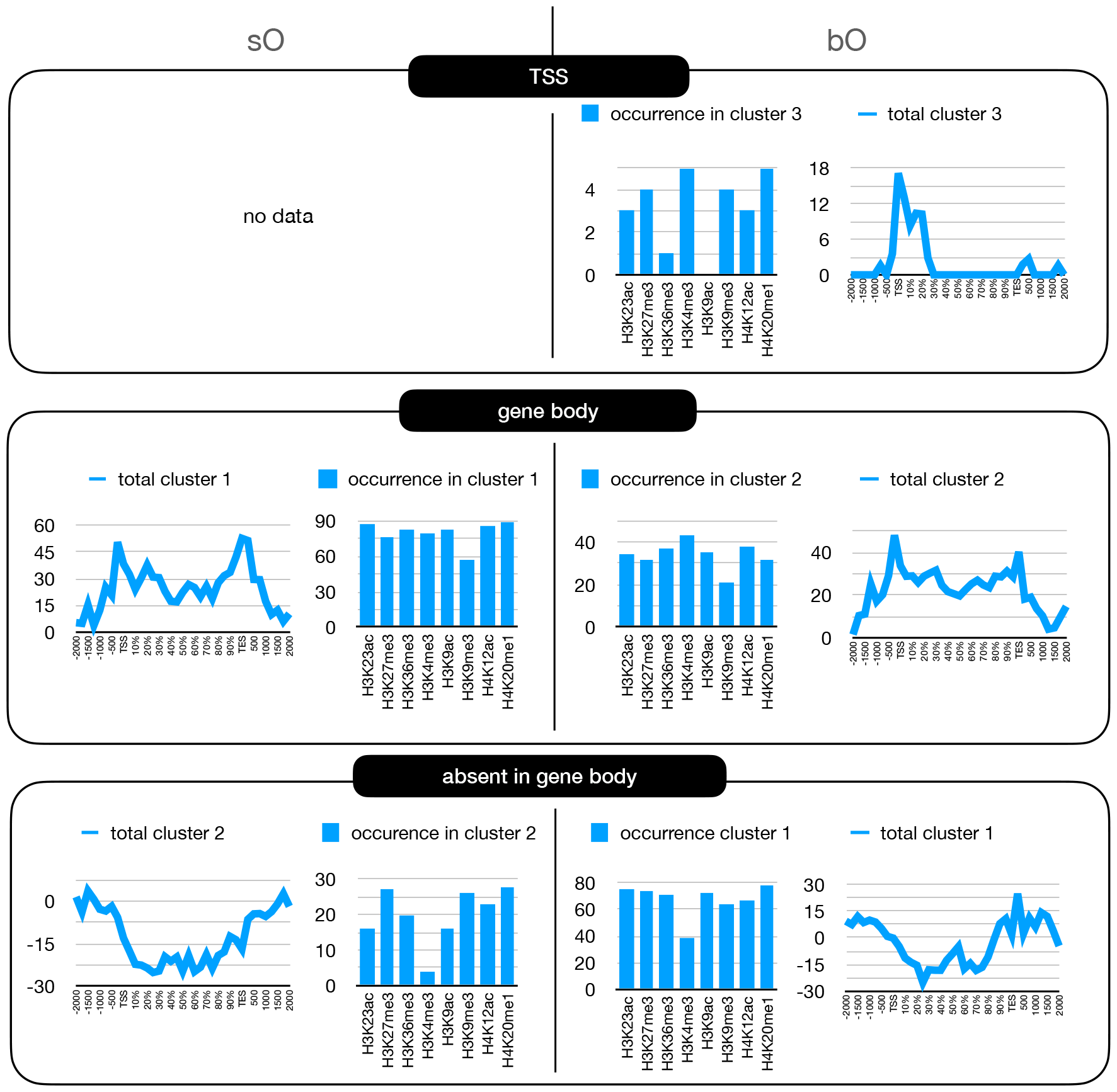
Metagene profiles of ovaries of single sex, pairing-unexperienced females (sO), and bisex, pairing-experienced females (bO). Axis as for figure 1a.

## Discussion

Our results reveal a complex landscape of chromatin modifications surrounding long non-coding RNA (lncRNA) genes in the gonads of *S. mansoni*, with a theoretical capacity to generate 256 distinct “chromatin colors” through combinations of eight histone modifications. Remarkably, in the gonads, we observed nearly the full spectrum of these combinations around lncRNA genes, yet these many modifications cluster into fewer than six categories based on the similarities of their metagene profiles. This clustering seems to be associated with distinct steps of gene transcription and represents a significantly reduced complexity compared to what might be expected by chance. Such findings suggest a functional role for histone modifications in as a regulatory mechanism that selectively employs certain histone modifications to orchestrate lncRNA transcription.

The discovery that the composition of chromatin colors varies between male and female gonads additionally indicates a sex-dependent difference. Furthermore, the pairing status of the worms obviously adds another layer of complexity to the regulation of lncRNA transcription. This sexual dimorphism and pairing-status specificity in chromatin modification patterns suggest that the regulation of lncRNA transcription is finely tuned to the physiological and developmental context of the parasite, especially with respect to the pairing status. The differential chromatin landscapes between males and females and between different pairing statuses imply that there is a strong epigenetic component in form of histone modifications that might play key roles in mediating sex-specific and pairing-dependent gene expression programs.

Our results suggest that the precise control of lncRNA transcription via specific patterns of histone modifications may represent a critical aspect of these regulatory functions. It is conceivable that aberrations in these chromatin modification patterns could lead to dysregulated lncRNA expression and contribute to developmental malfunction. Indeed, a recent functional study in *S. mansoni* identified lncRNA involvement in regulation cell proliferation in adults and gonads as well as reproduction including egg development (Silveira et al., 2023)

In conclusion, our study provides additional compelling evidence for the functional role of histone modifications in the control of lncRNA transcription in *S. mansoni* gonads, revealing the possibility of a regulatory system that is sensitive to the parasites′ developmental and physiological context. These findings open new avenues for understanding the mechanisms of gene regulation at the chromatin level and their implications for development, and physiology of the parasite, and probably also for the control of schistosomiasis, a NTD for which new treatment concepts are urgently needed (Bergquist et al., 2017; Silveira et al., 2022)

## Materials and Methods

### Life cycle maintenance of S. mansoni and ethics statement

To maintain the life cycle of the Liberian strain of *S. mansoni*, (Grevelding, 1995), we used *Biomphalaria glabrata* as intermediate host and hamster as final hosts in accordance with the European Convention for the Protection of Vertebrate Animals Used for Experimental and Other Scientific Purposes (ETS No 123; revised Appendix A). Animal experiments were approved by the Regional Council Giessen (V54-19 c 20/15 c GI 18/10).

### Organ extraction and N-ChIP

To generate worm populations of one sex, we performed monomiracidial infections of snails, in which clonal cercarial populations (single-sex cercariae) develop, and which we subsequently used for final host infections to obtain pairing-unexperienced females or males. Mixed-sex worm populations were obtained by polymiracidial snail infections.

Gonad isolation was performed by a detergent- and enzyme-based isolation procedure (Hahnel et al., 2013) using about 50 pairing-experienced females and males, and 100 pairing-unexperienced females, which are much smaller than paired females, as are their ovaries. For vitality check, 5-10 organs of each sample were stained 10 µl Trypan Blue (0.4%; Sigma) and checked by phase-contrast microscopy (Olympus IX 81 microscope).

N-ChIP was done as described before (Cosseau et al., 2009). Antibodies were purchased from commercial suppliers.

### Alignment and ChromstaR optimization

Uniquely aligned reads for N-ChIP with antiH3K4me3 on two biological replicates of pairing experienced ovaries (bF) and pairing unexperienced ovaries (sF) were downsampled to 2.5 Mio reads per sample and the resulting BAM files were used as input for peak calling in MACS2 (Effective genome size 350 Mb, building a model with lower mfold bound 5, upper mfold bound 50, band width for picking regions to compute fragment size 300, Peak detection based on qvalue, Minimum FDR (q-value) cutoff for peak detection 0.05), Peakranger 1.17 (Peak detection mode at low resolution, read extension length 300 bp), both with input BAM DU Sm-SsOv-INPUT. ChromstaR was evoked with default parameters and 250, 500, 750 and 1 000 bp bin size, with or without input.

Metagene profiles were generated with DeepTools for MACS and Peakranger results, and with plotEnrichment function of ChromstaR using 5 275 plus strand protein coding genes based on the Smv7 annotation (ftp://ftp.sanger.ac.uk/pub/project/pathogens/Schistosoma/mansoni/v7/annotation/Sm_v7.1.gff). Result files were visually inspected on IGV. We used version 7 of the genome assembly (ftp://ftp.sanger.ac.uk/pub/project/pathogens/Schistosoma/mansoni/v7/sequence/Smansoni_v7.fa).

### Metagene analysis around lncRNA genes and clustering by color

Coding sequences of lncRNA were obtained as BED file from (Vasconcelos et al., 2018). Metagene profiles were generated with plotEnrichment function of ChromstaR using 8,328 lncRNA coding genes on the plus strand. Metagene profiles over all combinations were exported from ChromstaR as individual TSV files. Hierarchical clustering was performed using the hclust function of R (figure 2).

**Fig. 2.**
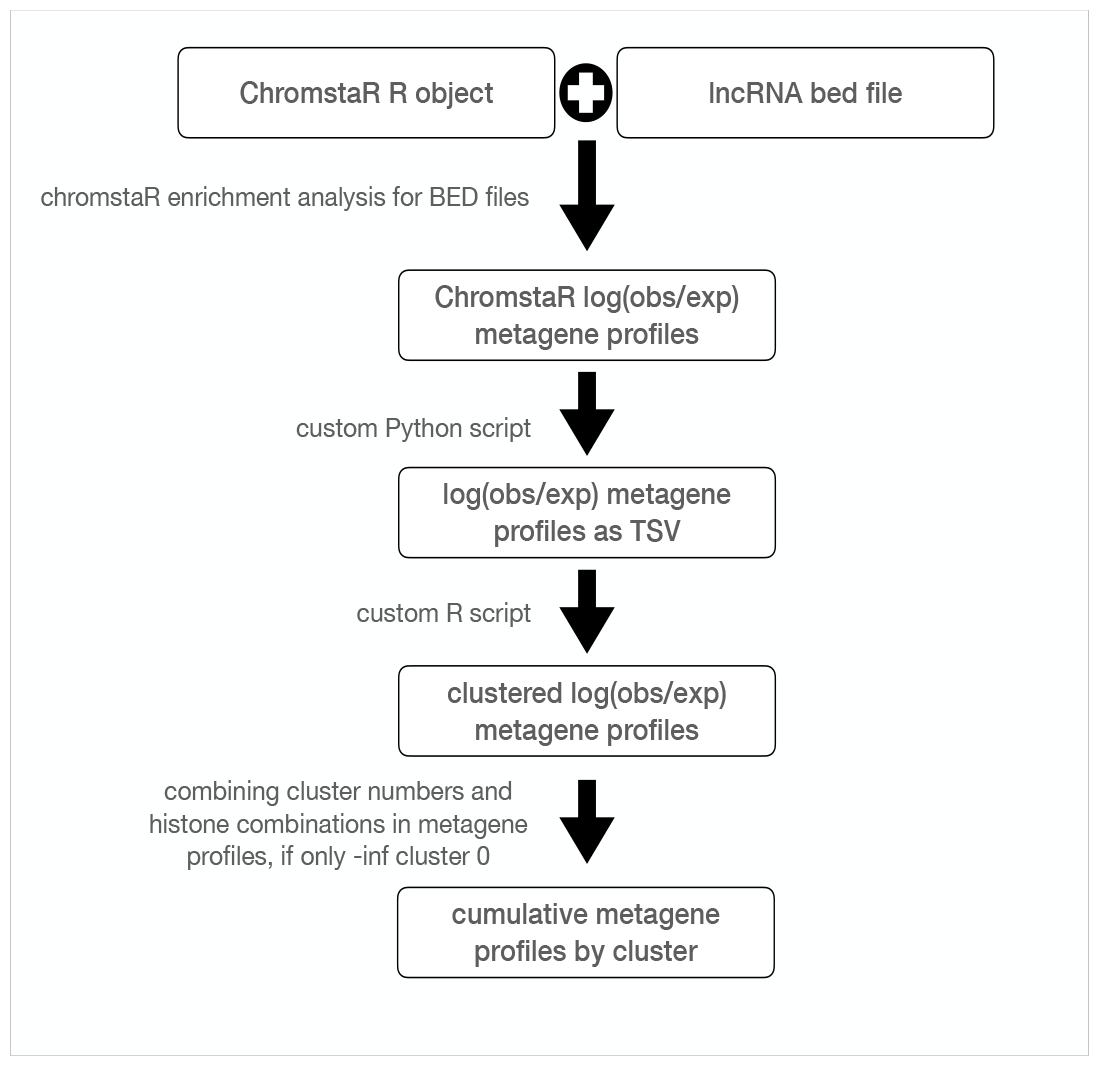
Schematic representation of the bioinformatics workflow used to perform clustering of metagene profiles.

## End Matter

### Author Contributions and Notes

C.G.G and C.G designed research, R.A and T.Q performed research; and all authors wrote the paper. The authors declare no conflict of interest. This article contains supporting information that can be requested from the authors.

## Acknowledgments

This study is set within the framework of the “Laboratoires d’Excellences (LABEX)” TULIP (ANR-10-LABX-41). With the support of the Wellcome Trust (107475/Z/15/Z) and the LabEx CeMEB, an ANR “Investissements d’avenir” program (ANR-10-LABX-04-01).

## Notes

### Competing Interest Statement

The authors have declared no competing interest.

